# Human Vascularized Bile Duct-on-a Chip: A multi-cellular micro-physiological system for studying Primary Sclerosing Cholangitis

**DOI:** 10.1101/2023.03.02.530888

**Authors:** Yu Du, Kapish Gupta, Orith Waisbourd-Zinman, Adi Har-Zahav, Carol J. Soroka, James L. Boyer, Jessica Llewellyn, Chengyang Liu, Ali Naji, William J. Polacheck, Rebecca G. Wells

## Abstract

Primary sclerosing cholangitis (PSC) is a chronic cholestatic liver disease in which the bile ducts of the liver become inflamed and scarred. Scarred bile ducts eventually narrow and obstruct and can cause additional liver pathology including liver failure, repeated infections, and tumors. The pathogenesis of PSC remains largely unknown, partly due to difficulty in obtaining cholangiocytes and partly due to a paucity of *in vitro* models that capture the various factors contributing to disease progression. Here we report the development of a human vascularized bile duct-on-a-chip that models blood vessels and bile ducts structurally and functionally in three dimensions and includes cholangiocytes derived from control and PSC patient tissue and bile. The flow of blood and bile was modeled by perfusion of cell-lined channels, and cholangiocytes and endothelial cells displayed differential responses to perfusion. Normal and PSC cholangiocytes polarized normally, formed mature tight junctions and displayed similar permeability, comparable to *ex vivo* measurements. The model with PSC cholangiocytes, however, became more inflammatory than the normal under the stimulation of IL-17A, which induced PBMC and differentiated Th17 cells in the vascular channel to transmigrate more through the endothelial layer of the vascular compartment. In sum, this human vascularized bile duct-on-a-chip recapitulated the vascular-biliary interface structurally and functionally and represents a novel multicellular platform to study inflammatory and fibrotic cholangiopathies such as PSC.

## Introduction

Primary sclerosing cholangitis (PSC) is a rare, chronic cholestatic liver disease characterized by persistent and progressive inflammation and fibrosis around the bile ducts (1, 2). Although the pathogenesis of PSC remains largely unknown, evidence suggests it is a multifactorial cholangiopathy involving cells at the vascular-biliary interface including cholangiocytes, immune cells, endothelial cells, and mesenchymal cells (3–5). Animal models have been developed to model PSC, including Mdr2-/- mice, lithocholic acid-fed mice and TGR5-/- mice (6–8); however, none recapitulate all the aspects of the disease. Mouse models also suffer from potential confounding from inter-species variation, and dissecting the complex interactions between various cell types during disease progression can be challenging *in vivo*.

Organoids derived from human cells have demonstrated the advantage of propagating primary cells that maintain their original phenotype (9) and have been developed to model cholangiopathies including biliary atresia, Alagille Syndrome, and PSC (9–11). While these avoid the problems associated with rodent systems, they have a non-tubular 3D structure, lack flow and are hard to model as multicellular systems. Microfluidic organs-on-chips, another kind of model system, offer the potential to incorporate 3D structure, multiple cell types, and mechanical stimuli (12). We previously reported the development of a bile duct-on-chip that incorporates a tubular duct and exhibits normal cholangiocyte apical-basal polarity, barrier function and physiological responses to applied flow (13). Organs-on-chips, however, face the challenges of cell source and cellular fidelity; this is particularly problematic for a rare disease like PSC. Integrating these two methods – organoids and organs-on-chips – offers a way to bypass the limitations of each and provide an *in vitro* approach capable of advancing the study of cholangiopathies (14).

We report here the development of a novel multicellular *in vitro* platform, a human vascularized bile duct-on-a-chip (VBDOC) incorporating cholangiocytes from control and PSC patients with human fibroblasts and endothelial cells. We used the VBDOC to recapitulate the barrier function of blood vessels and bile ducts, test the response to mechanical force, and study cytokine secretion and immune cell transmigration in response to inflammatory stimuli. The human VBDOC represents a novel multicellular *in vitro* platform to study the pathophysiology of the biliary system using cholangiocytes from a variety of sources.

## Materials and Methods

### Isolation of primary human cholangiocytes

Primary extrahepatic bile ducts were obtained from previously healthy deceased organ donors (one 51 year old female, one 66 year old male) as part of the Human Pancreas Procurement and Analysis Program (HPPAP), which was granted IRB exemption (protocol 826489); cholangiocytes were isolated as described previously (15). PSC cholangiocytes from tissue were obtained from a surgical specimen (from a 40 year old male) and isolated similarly (15), with ethics approval was granted by Beilinson Medical Ethics Committee (approval number: 0072-19-RMC). Bile-derived PSC cholangiocytes were isolated as described (11, 16) with IRB exemption granted (protocol 2000020797).

### Generation and culture of organoids

For organoid culture, primary cholangiocytes were centrifuged at 444g for 4 min and resuspended in a mixture of 66% Matrigel (BD Biosciences, 356237) and 33% complete cholangiocyte culture medium (CCCM: William’s E medium (Gibco, Life Technologies) supplemented with 10 mM nicotinamide (Sigma-Aldrich), 17 mM sodium bicarbonate (Sigma-Aldrich), 0.2 mM 2-phospho-l-ascorbic acid trisodium salt (Sigma-Aldrich), 6.3 mM sodium pyruvate (Invitrogen), 14 mM glucose (Sigma-Aldrich), 20 mM HEPES (Invitrogen), ITS + premix (BD Biosciences), 0.1 μM dexamethasone (R&D Systems), 2 mM Glutamax (Invitrogen), 100 U/ml penicillin and 100 μg/ml streptomycin, 20 ng/ml EGF (R&D Systems), 500 ng/ml R-spondin (R&D Systems) and 100 ng/ml DKK-1 (R&D Systems)) with 10 µM Y27632 (Sigma-Aldrich). The cell suspension was plated in a 24-well plate format at 300 μl/well (~1×10^5^ cells), and plates were then incubated at 37°C for 30 min until the Matrigel solidified. Subsequently, 1 ml of CCCM with 10 µM Y27632 was added to each well and the culture medium was switched to CCCM after 2 days. The culture medium was changed every 2 days.

### Culture and transfection of human gall bladder fibroblasts

Primary human gallbladder fibroblasts were purchased from ScienCell (5430), and cultured using the fibroblast growth medium kit from Lonza (CC-3132). GFP-labelled cells were generated by infecting them with GFP lentivirus (Addgene 17448).

### Culture of HUVECs

HUVECs were cultured using the endothelial growth medium (EGM) kit from Lonza (CC-2519, CC-3202) according to the supplier’s instructions. HUVECs under passage 7 were used for all experiments.

### Fabrication of the VBDOC

Microfluidic molds were fabricated using high-resolution 3D printing by Proto Labs, Inc. (Maple Plain, MN). Polydimethylsiloxane (PDMS, Sylgard 184, Dow-Corning, Midland, MI) devices were cast by pouring the PDMS mixture of base and curing agent (weight ratio 10:1) into the mold, followed by degassing and full cure at 75°C for 1.5 h. Cured PDMS gel was cut into bricks of equal size followed by punching ports and cleaning with tape. The PDMS was bonded to the coverslip by 45 s of plasma treatment. The collagen gel chamber was treated with 0.01% (v/v) poly-L-lysine for 1 h and 0.5% (v/v) glutaraldehyde for 20 min to promote collagen adhesion. After the devices were washed overnight in water and for 30 min in 70% ethanol, steel acupuncture needles (200 μm diameter, Seirin, Kyoto, Japan) were inserted from opposite directions and the devices were then sterilized under UV for 20 min in neutralized collagen solution (2.5 mg/ml, pH=7.0), which was prepared by mixing rat tail type 1 collagen (Thermo Fisher Scientific), 10X DMEM medium, 10 mM HEPES, 1 M NaOH and NaHCO_3_ (0.035% w/v) on ice. Fibroblasts in culture dishes were then trypsinized, washed and resuspended with the neutralized collagen solution at a density of 1.5×10^5^/ml. The fibroblast/collagen mixture was injected through the side ports to fill the collagen gel chamber and the collagen was allowed to gel by incubating the inverted devices at 37°C for 20 min. Fibroblast growth medium was added to each port and incubated overnight, then needles were removed to yield channels. A suspension of 5×10^5^/ml of cholangiocytes was introduced to the reservoir ports connected to the cholangiocyte channel. Cells were allowed to adhere to the top surface of the channel for 5 min; devices were then flipped to allow cells to adhere to the bottom surface of the channel for 5 min. (Times were determined empirically to yield an even cell coating of the channel and sufficient adhesion to permit rinsing the channel.) Nonadherent cells were removed by rinsing with cell-culture medium, and the devices were filled with fresh CCCM with 10 µM Y27632. Devices were maintained at 37°C (5% CO_2_) on a rocker at 5 rpm for 2 d. The medium was switched to CCCM with medium changes every 2 days until the development of confluent monolayers. When the cholangiocyte monolayer was just confluent, HUVECs in culture dishes were trypsinized, washed and resuspended with EGM at a density of 5×10^5^/ml. The cell suspension was introduced to reservoir ports connected to the endothelial channel. Cells were allowed to adhere to the top surface of the channel for 2 min; devices were then flipped to allow cells to adhere to the bottom surface of the channel for 2 min. (Times were again determined empirically.) Nonadherent cells were removed by rinsing with EGM. The reservoir ports connected to the endothelial channel were filled with EGM while the reservoir ports connected to the cholangiocyte channel were filled with CCCM. Devices were maintained at 37°C (5% CO_2_) on a rocker at 5 rpm (static groups were maintained at 37°C (5% CO_2_) on a flat incubator shelf) until the development of confluent endothelial monolayers and compact cholangiocyte monolayers(13).

### Permeability measurements

To measure the permeability of the cholangiocyte and endothelial cell monolayers in the channels, fluorescent dextran (70, 10, and 4 kDa, labeled with fluorescein isothiocyanate (FITC); Sigma-Aldrich) in phosphate-buffered saline (PBS) was introduced into the channels at a concentration of 20 μg/ml. Diffusion of the dextran was imaged in real time with an EVOS FL Auto 2 Imaging System (Thermo Fisher Scientific) at 10× magnification. The diffusive permeability coefficient was calculated by measuring the flux of dextran into the collagen gel and fitting the resulting diffusion profiles to a dynamic mass conservation equation, as described (13, 17).

### Immunostaining

Cholangiocyte and endothelial cell monolayers in the device were fixed with 4% paraformaldehyde (PFA) at 37°C for 20 min with rocking. Cells were rinsed 3× with PBS and permeabilized with 0.1% Triton X-100 for 3 days, then blocked with 2% bovine serum albumin (BSA; Sigma-Aldrich) in PBS at 4°C overnight with rocking. Primary antibodies, together with 4′,6-diamidino-2-phenylindole (DAPI; Thermo Fisher Scientific) diluted in 2% BSA in PBS, were incubated overnight at 4°C and rinsed 3× with PBS for 5 min each with rocking, followed by an overnight rinse. Secondary antibodies were diluted in 2% BSA in PBS, incubated overnight at 4°C, and rinsed 3× with PBS for 5 min each on a rocker. Primary antibodies and the concentrations used are listed in Supporting Table S1. Cyanine (Cy)3- and Cy5-conjugated secondary antibodies were used at 1:400 (Vector Laboratories, Burlingame, CA).

### Cytokine array

Cholangiocyte monolayers were cultured in CCCM with or without IL-17A for 48 h, after which the media in all four reservoirs was collected and analyzed with the Human Cytokine Antibody Array (Abcam, Cambridge, UK). The full list of cytokines tested in the kit is found in supporting Table S2.

### Rhodamine 123 transport assay

Rhodamine 123 transport assays were performed in the VBDOC with only cholangiocytes in the device. 5 µM Rhodamine 123 in the CCCM was perfused to the basal side through the endothelial channel (without endothelial cells) and normal CCCM was added to the cholangiocyte channel after blocking the cholangiocyte channel ends with vacuum grease – this ensured that there was no Rhodamine 123 in the lumen at the start of the assay. After incubating for 2 h at 37°C, the basal side was washed 3 times with PBS through endothelial channel.

In the inhibition assay, cholangiocytes were incubated with10 µM Verapamil (Sigma-Aldrich) in CCCM at 37°C for 30 min, followed by incubation with Rhodamine 123 as described above. Images were acquired using a SCTR Leica confocal microscope and Leica application suite (LAS X; Leica, Buffalo Grove, IL). Fluorescence intensity was measured across the cholangiocyte channel.

### *In vitro* Th17 differentiation

CD4 T cells were provided by the Human Immune Core at the University of Pennsylvania. They were cultured on anti-CD3 (5 µg/ml in PBS, overnight at 4°C)-coated plates in complete RPMI (RPMI1640+Glutamax medium, supplemented with 10% FCS and 1% P/S). For Th17 differentiation, CD4 T cells were plated at a density of 2X10^6^ cells per well (24 well plates) with 50 ng/ml of human IL23 and IL1b in 500 µl complete RPMI. On day 4, medium and cytokine mix were replaced. On day 7, cells were collected for transmigration assays and Th17 levels were analyzed by flow cytometry.

### Transmigration assay

After cholangiocytes and endothelial cells reached confluence in their individual channels in the VBDOC, the cholangiocytes were treated with 50 ng/ml IL-17A in culture medium for 48 h; for the last 4 h, the endothelial cells were also treated with 1 µg/ml LPS. PBMCs provided by the Human Immune Core of University of Pennsylvania or differentiated CD4T cells were fluorescently labeled in RPMI with 5 µM Celltracker Green CMFDA for 30 mins at 37°C and resuspended in 2X10^6^/ml EGM after washing with EGM. 110 µl of cell suspension was added to one side of the reservoir port, 90 µl was added to the other side of the reservoir port, and the devices were placed in the incubator overnight. Images of transmigrated cells in the VBDOC were taken by confocal z-stack scanning.

### Statistical analysis

Statistical significance was assessed using one-way ANOVA. p<0.05 was regarded as statistically significant and calculated with Prism 7 (GraphPad Software, La Jolla, CA). All data are presented as mean ± standard deviation (SD). Sample size is indicated in the corresponding figure legends.

## Results

### Fabrication of the vascularized bile duct-on-a-chip

In order to recapitulate the vascular-biliary-mesenchymal interface, we designed a two-channel microfluidic device modified from a previously-described model of sprouting angiogenesis(18). To generate a vascular channel and bile duct with an intervening submucosa-like region, a collagen solution containing human gallbladder fibroblasts was injected through the side ports of a PDMS device (Fig. 1A,B) and gelled around two parallel needles (200 µm) inserted through the chamber in the middle of the device. The two channels formed after removing the needles were connected to separate pairs of reservoirs. Cholangiocytes derived from organoid culture were seeded into one channel and self-organized into a confluent and then compact epithelial monolayer after about two weeks. When the cholangiocyte monolayer became confluent, endothelial cells were seeded in the other channel, developing into a confluent monolayer after 3 days (Fig. 1D). The vascular vessel, bile duct and surrounding mesenchymal cells in the VBDOC (Fig. 1) maintained structural integrity for at least a week.

**Figure 1.**
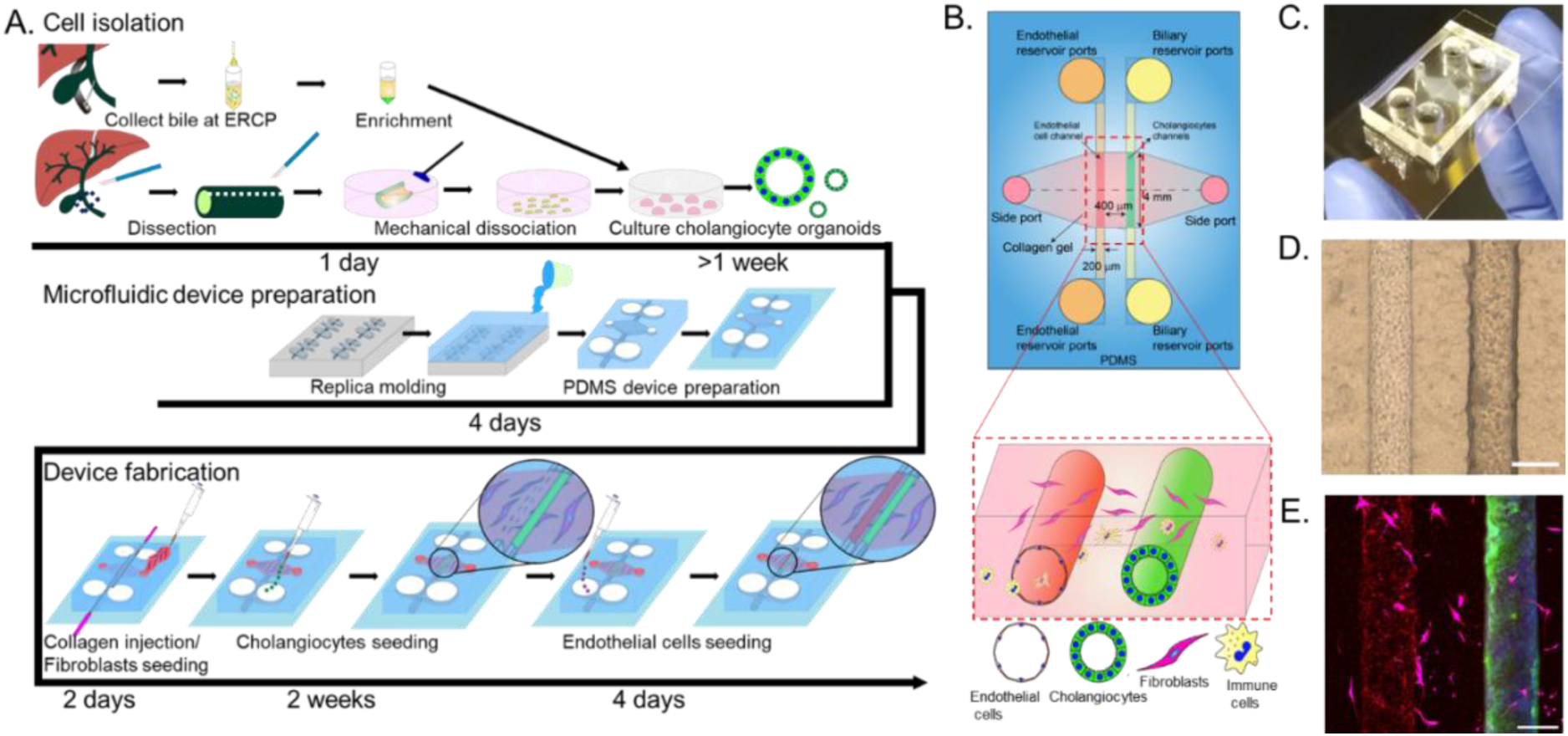
Fabrication of a human VBDOC using organoid-derived cholangiocytes. A. Flow chart showing an overview of the process of cell isolation, microfluidic device preparation and device seeding. B. Schematic showing top and cross-sectional views of the VBDOC. C. Image of a typical VBDOC, top view. D. Representative bright field image of the endothelial (left) and biliary (right) channels with fibroblasts in the surrounding matrix. E. Representative immunofluorescence images of endothelial cells (VE-cadherin, red), fibroblasts (transfected GFP, magenta) and cholangiocytes (K19, green) in the VBDOC. Nuclei shown by DAPI staining (blue). Images are representative of at least three independently-constructed and seeded devices for each condition. Scale bars: 200 μm.

### Characterization of the normal and PSC organoids-derived cholangiocytes in the device

We generated cholangiocyte-lined channels successfully using cholangiocytes dissociated from tissue-derived organoids from both control and PSC patients, then characterized them with immunofluorescence staining. Cells maintained expression of the cholangiocyte markers K7, K19, GGT and EPCAM (Figs. 1E, 2). Apical F-actin and primary cilia with basal collagen IV and laminin (produced by the cholangiocytes themselves, since there was none coating the channel) demonstrated that cholangiocytes are polarized in the device (Fig. 3). To broaden the cell source, especially from PSC patients, we also seeded cholangiocytes dissociated from bile-derived organoids from PSC patients into the devices (11). The bile-derived cholangiocytes, like those from tissue, survived and generated confluent monolayers in the devices. Immunofluorescence staining for K7, K19, GGT, EPCAM, F-actin, collagen IV, laminin, and acetylated α-tubulin (for primary cilia) showed similar expression of biliary markers and polarity with less continuous laminin and less acetylated α-tubulin compared to the tissue-derived cholangiocytes in the devices (Fig. 2, 3).

**Figure 2.**
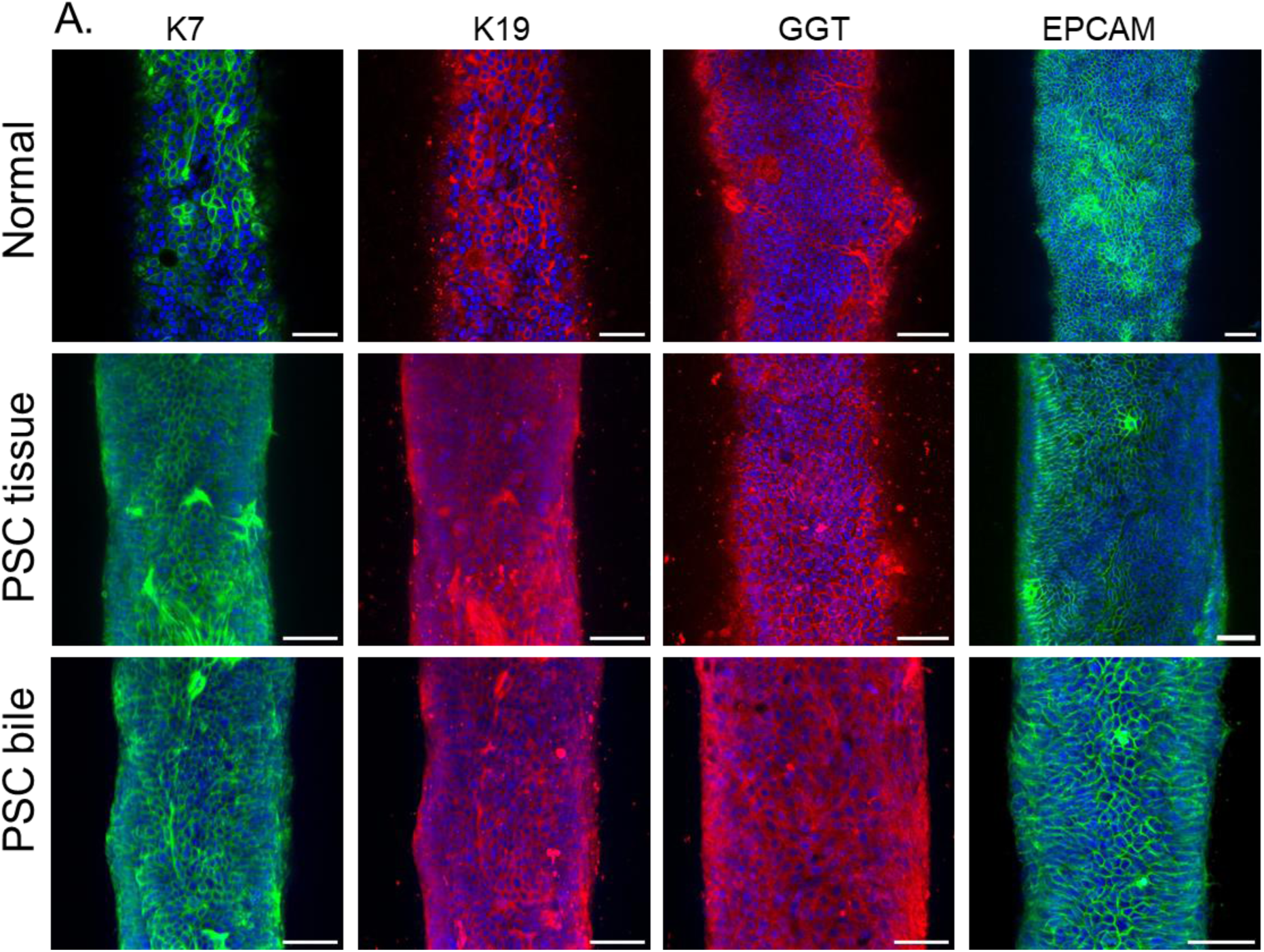
Characterization of cholangiocytes in channels. Channels lined with normal cholangiocytes derived from control tissue (top row), PSC cholangiocytes from tissue (middle) and PSC cholangiocytes from bile (bottom). Representative confocal images of cholangiocytes in the device stained with antibodies against K7, K19, GGT and EPCAM. Nuclei shown by DAPI staining (blue). Images are representative of at least three independently-constructed and seeded devices for each condition. Scale bars: 50 μm.

**Figure 3.**
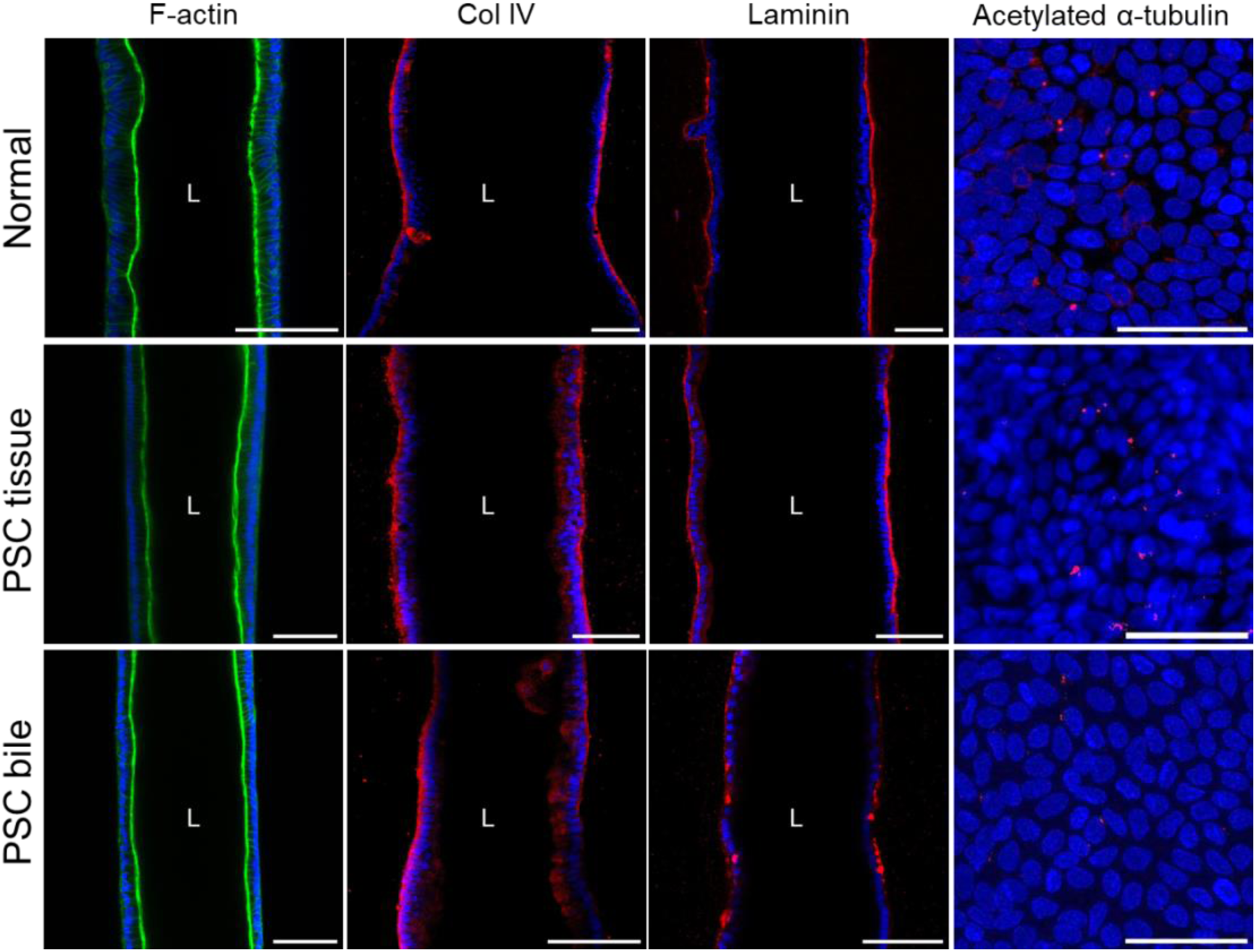
Characterization of cholangiocyte polarity in the VBDOC. Channels lined with normal cholangiocytes derived from control tissue (top row), PSC cholangiocytes from tissue (middle) and PSC cholangiocytes from bile (bottom). Immunofluorescence images of the channel stained with antibodies against F-actin, Collagen IV, Laminin (middle cross-sectional view), and acetylated α-tubulin (top projection view). Nuclei shown by DAPI staining (blue). Images are representative of at least three independently-constructed and seeded devices for each condition. Scale bars: 100 μm, 50 μm in last column.

### Bile salt transporter expression and activity in the device

We compared transporter expression between normal (Fig. 4A) and PSC cholangiocytes from tissue (Fig. 4C) and bile (Fig. S1) in the channel by immunostaining for the apical sodium-dependent bile acid transporter (SLC10A2; also known as ASBT), the channel CFTR and the basal secretin receptor (SCTR). Transporter activity was interrogated by examining the ability of cholangiocytes to transport rhodamine 123 into the luminal side of the cholangiocyte channel (Fig. 4B, D). Confocal microscopy showed that the cholangiocytes lining the channel took up rhodamine 123 and secreted it into the lumen, thereby confirming active transport through MDR1. This luminal extrusion of rhodamine 123 was inhibited by the MDR1 inhibitor verapamil. Similar uptake and efflux were seen in the channels lined with cholangiocytes from normal and PSC organoids. Our data therefore demonstrate that both normal and PSC cholangiocytes from organoids maintain their functional properties in the devices.

**Figure 4.**
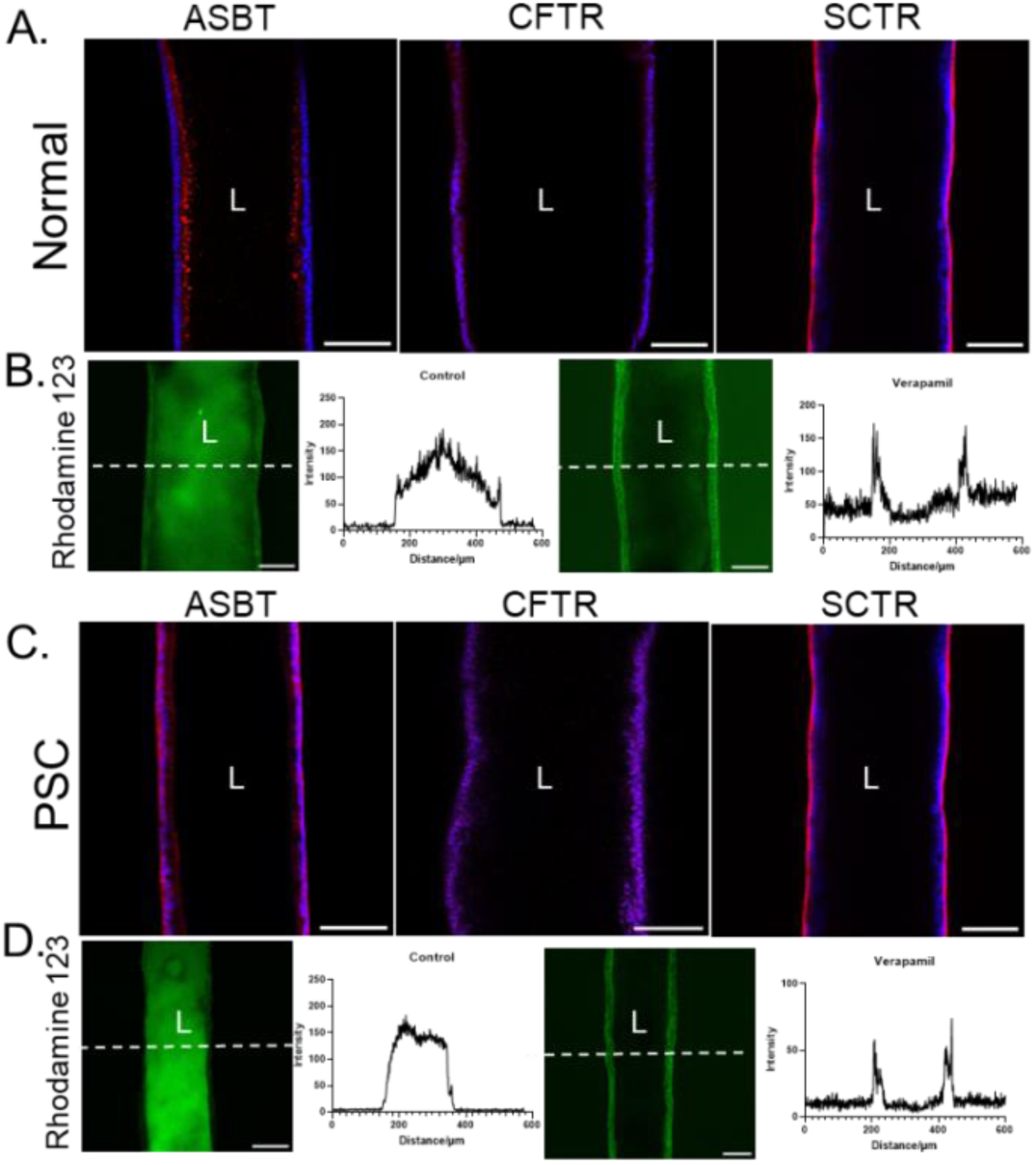
Expression and functional characterization of bile salt transporters in cholangiocytes lining the biliary channel. A,C) Immunofluorescence images of channels lined with normal (A) and PSC tissue (C) cholangiocytes stained with antibodies against ASBT, CFTR and SCTR (middle cross-sectional view). Nuclei shown by DAPI staining (blue). Scale bars: 100 μm. B,D) Secretion of fluorescent rhodamine 123, a substrate of MDR1, into the lumen of the cholangiocyte channel (left), with inhibition by the MDR1 inhibitor verapamil (right). Graphs depict the fluorescence intensity along the dotted white lines in the images. Images are representative of at least three independently-constructed devices for each condition.

### Barrier function of normal and PSC cholangiocytes in the VBDOC

We next examined barrier function in the cholangiocyte channel. By immunofluorescence staining, there was no difference in expression of E-cadherin, ZO-1, occludin or claudin-1 at cell-cell junctions between normal and PSC cholangiocytes in the devices (Fig. 5A). To compare the barrier function of the normal- and PSC-cell seeded bile duct channels, we perfused the cholangiocyte channels with FITC-dextran ranging in size from 4 to 70 kDa. There was no obvious leakage of fluorescent dextran of any size into the collagen matrix even after 10 min (Fig. 5B). Quantification of permeability showed no significant difference between normal and PSC cholangiocyte channels (Fig. 5C).

**Figure 5.**
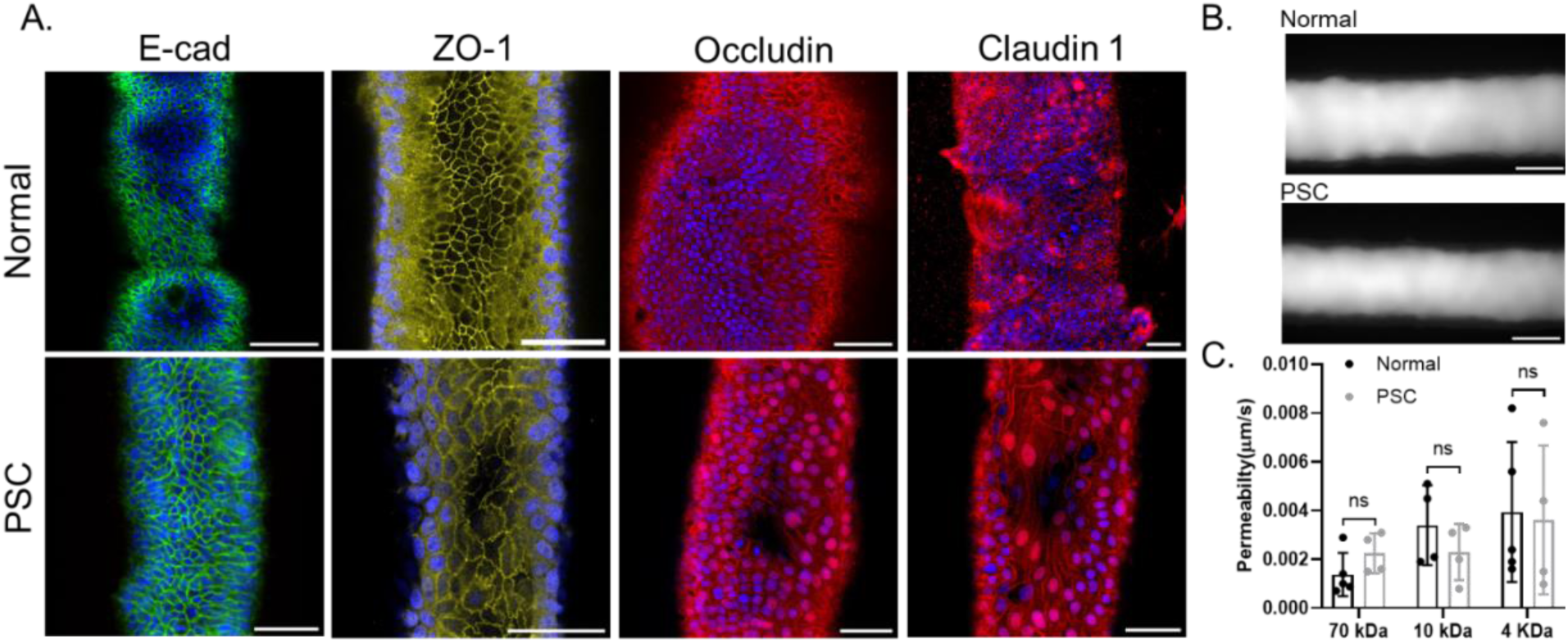
Barrier function of normal and PSC cholangiocytes. A. Immunofluorescence images of cholangiocyte channels lined with control (upper row) and PSC (lower row) tissue cholangiocytes stained with antibodies against E-cadherin, ZO-1, occludin and claudin-1. Nuclei shown by DAPI staining (blue). Scale bars: 50 μm. Images are representative of at least three independently-constructed devices for each condition. B. Representative image of FITC-dextran (70 kDa) in the channel, imaged after 10 min. Scale bar, 200 μm. C. Quantification of permeability of the control and PSC cholangiocyte-lined channels to FITC-dextran (70 kDa, 10 kDa and 4 kDa), n=4-5 devices, each device tested sequentially with FITC-dextran from 70 to 4 kDa. All data are presented as mean ± SD, *P<0.05.

### Different mechanosensitivity of cholangiocytes and endothelial cells to shear flow

In order to define the flow sensitivity of cholangiocytes compared to endothelial cells, the different cells were co-cultured in neighboring channels on a rocker. The rocking causes oscillating flow through the channels (17, 19), enabling us to examine the responses of cholangiocytes and endothelial cells to shear flow. We used F-actin and VE-cadherin immunostaining to assay for morphological changes. Cholangiocytes maintained a similar cuboidal shape and random alignment under both static and flow conditions. Endothelial cells, however, became elongated under flow and aligned with the direction of flow (Fig. 6A, B). Because barrier function is crucial for vascular and biliary function, we compared the permeability under static and flow condition for both types of cells (Fig. 6C, D). Consistent with data reported previously, flow promotes the establishment of a functional vascular barrier in the endothelial channel (17), as shown by decreased permeability. In contrast, the permeability of cholangiocyte monolayers remained constant with and without shear flow, with permeability comparable to ex vivo measurements (Fig. 6C, D) (20, 21).

**Figure 6.**
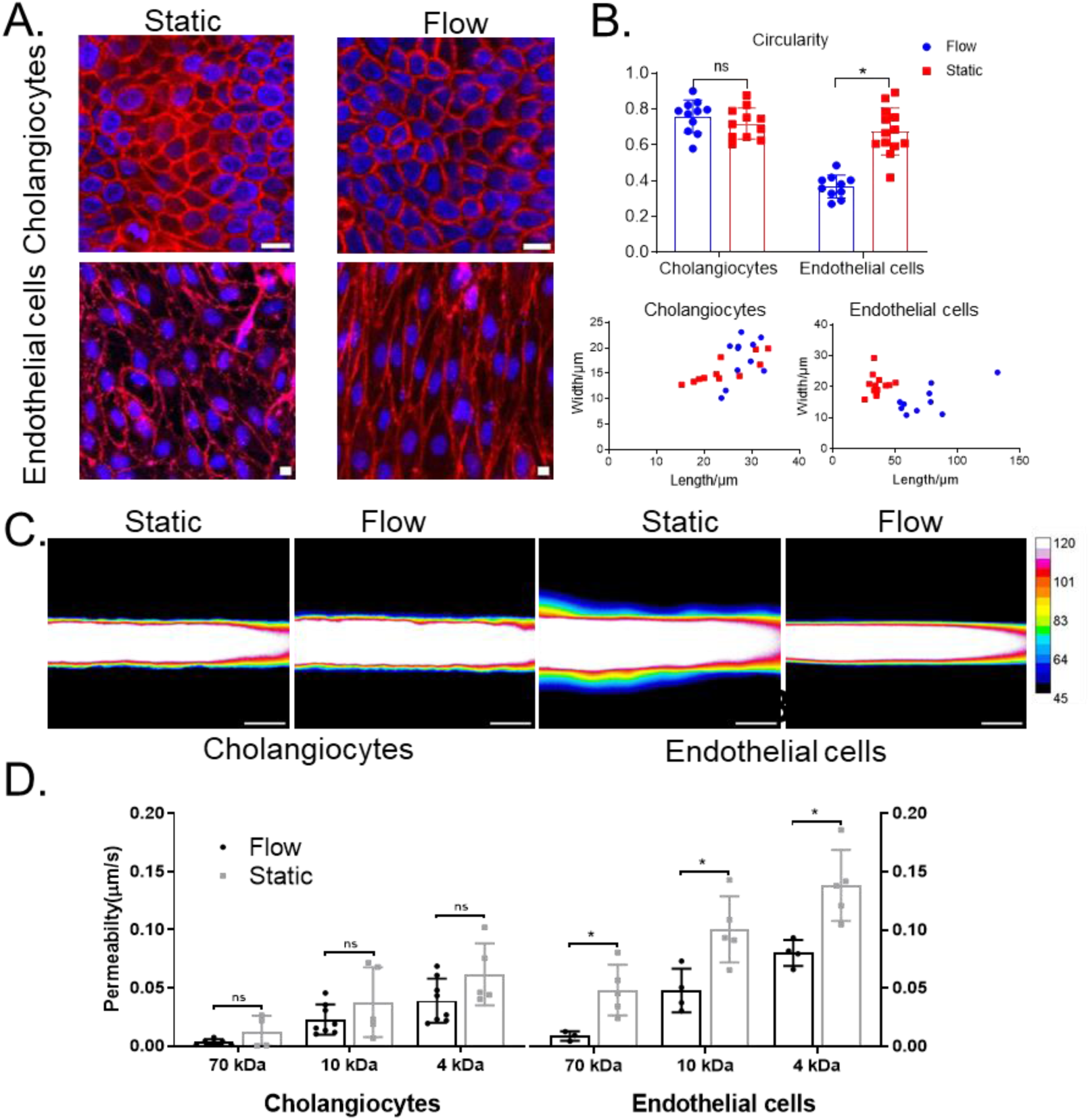
Morphological and functional responses of endothelial cells and control cholangiocytes to luminal flow. A. Representative confocal images of cholangiocytes stained with F-actin and endothelial cells stained with VE-cadherin under static or flow conditions in the VBDOC. Nuclei shown by DAPI staining (blue). Images are representative of at least three independently-constructed devices for each condition. Scale bars: 100 μm. B. Graphs quantifying length, width (lower panel) and circularity (upper panel) in endothelial cells and cholangiocytes under static or flow conditions in the VBDOC. C. Representative pseudo-color images of FITC-dextran (70 kDa) diffusion after 2 min in the cholangiocyte and endothelial channels of the VBDOC under static and flow conditions. Scale bar: 200 μm. D. Permeability of the cholangiocyte channel and endothelial channel to FITC-dextran (70, 10, 4 kDa) under static and flow conditions, n=4-8.

### Cytokine profile of normal and PSC cholangiocytes in the VBDOC with IL-17A stimulation

Cholangiocytes can react to inflammation and liver damage by acquiring a reactive inflammatory phenotype in which the ductular cells secrete proinflammatory and profibrotic chemokines and cytokines (3, 22). We tested whether control and PSC tissue cholangiocytes in the devices could be stimulated to secrete chemokines. Based on reports that IL-17 can stimulate biliary epithelial cells to secret CCL20 (11, 23), we incubated cholangiocytes with IL-17A (50 ng/ml for 48 h) and measured the cytokines secreted into the culture supernatant. Normal and PSC cholangiocytes expressed similar amounts of IL-8 and angiogenin with and without IL-17A stimulation, but demonstrated increased expression of CCL20, CXCL5, CXCL1, uPAR, and GRO α/β/γ. In response to IL-17A, normal cholangiocytes also increased expression of CXCL6 and CXCL7, while PSC cholangiocytes increased expression of TIMP-1 (Fig. 7). (PSC cholangiocytes from bile-derived organoids exhibited a similar inflammatory profile as PSC cholangiocytes from tissue-derived organoids, Figure S2). Cholangiocytes in the devices can therefore be stimulated to react in an inflammatory manner, including secreting CCL20, which has been studied previously (11, 23).

**Figure 7.**
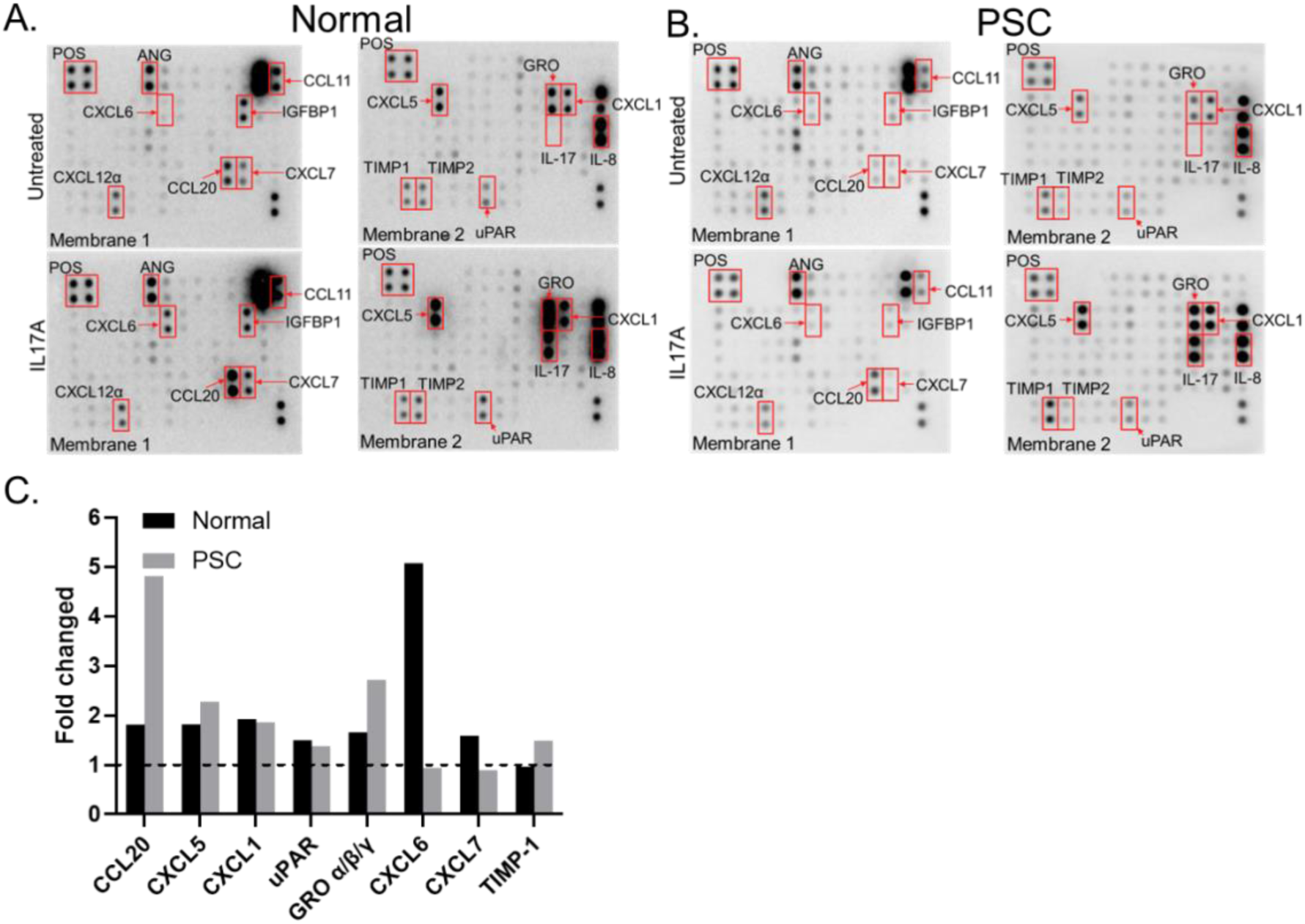
Cytokine profile of normal and PSC cholangiocytes in the VBDOC after stimulation with IL-17A. A, B. Human cytokine array of untreated (upper panels) and IL-17A treated (lower panels) control (A) and PSC (B) cholangiocytes in the biliary channel. C. Quantification of the blots, showing fold change in protein expression normalized to untreated samples.

### Immune responses in the VBDOC in response to IL-17A stimulation

It has been postulated that certain chemokines and cytokines, especially CCL20, attract T helper 17 (Th17) lymphocytes to damaged bile ducts in various liver diseases (23, 24). Bacterial infections are frequently detected around the portal tracts of PSC patients and elevated serum endotoxin can increase the expression of adhesion molecules such as VCAM-1 and ICAM-1 (24–26). To recapitulate inflammatory responses in PSC, we stimulated PSC cholangiocytes with IL-17A (50 ng/ml for 48 h) to generate an inflammatory phenotype. Simultaneously, we stimulated control endothelial cells in the same devices with LPS (1µg/ml for 4 h) to promote immune cell adhesion. LPS stimulation, as shown by immunofluorescence staining, increased the expression of ICAM-1 and VCAM-1 on endothelial cells (Fig. S3). To investigate whether IL-17A-stimulated PSC cholangiocytes can attract Th17 cells, CD4 T cells were isolated from human peripheral blood and differentiated *in vitro* (Fig. S4), as previously described (27, 28). Fluorescent-labeled human PBMC or differentiated CD4 T cells were then perfused into the endothelial channels and incubated overnight. PBMC and differentiated CD4 T cells adhered to endothelial cells and transmigrated into the surrounding matrix in all channels equally, regardless of LPS stimulation (Fig. 8). The number of transmigrated PBMC and differentiated CD4 T cells did not increase significantly under LPS stimulation but, notably, transmigration increased 18.6-fold (PBMC) and 3.5-fold (differentiated CD4 T cells) when cholangiocytes in the second channel had been stimulated with IL-17A. These findings demonstrated that the VBDOC can successfully model immune cell recruitment.

**Figure 8.**
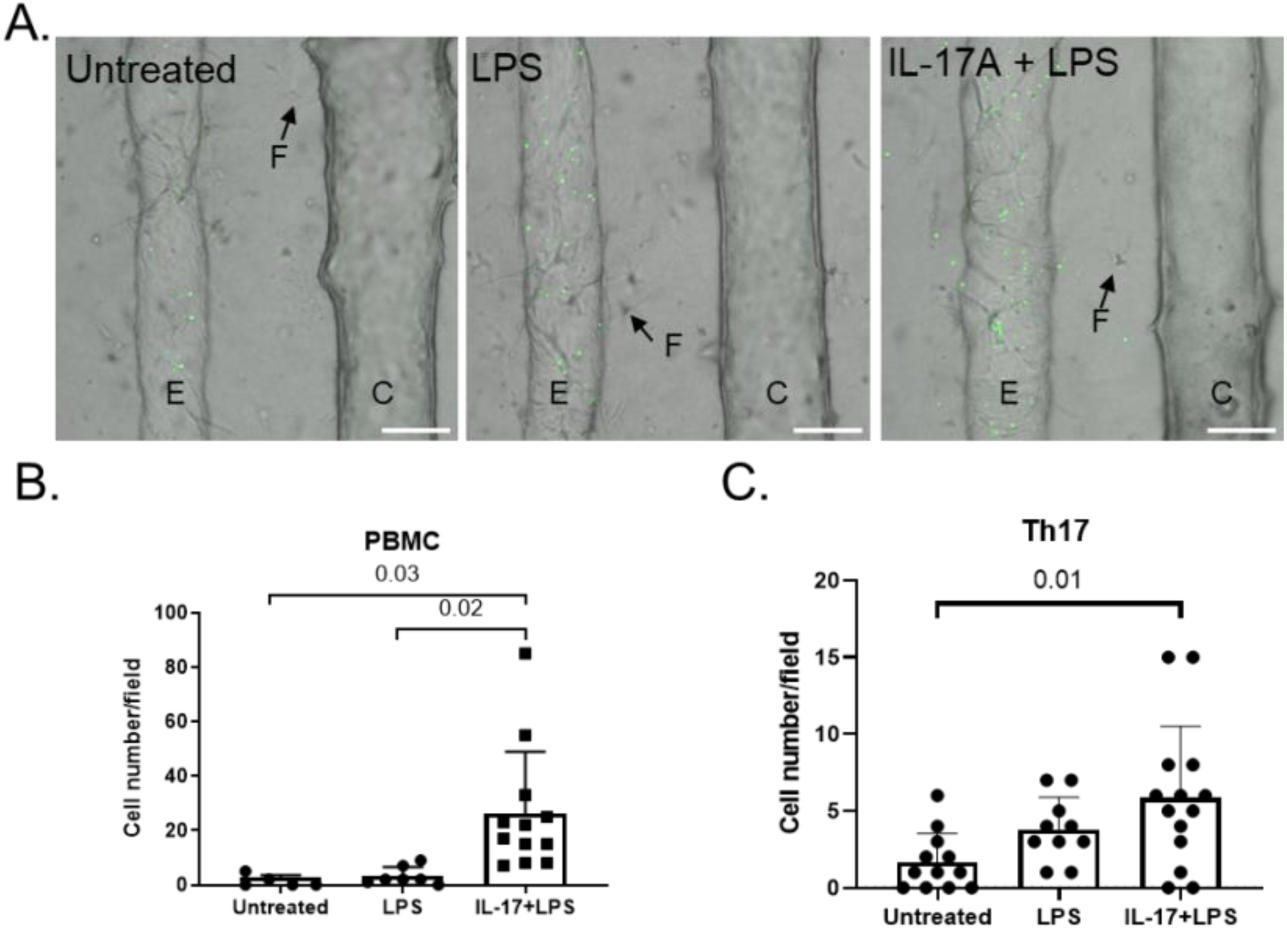
Immune cell transmigration in the VBDOC. A. Representative images of transmigration of cell tracker-labeled (green) immune cells through the endothelium when treated with vehicle, LPS, or LPS combined with cholangiocyte IL-17A stimulation. E: endothelial cells, C: cholangiocytes, F: fibroblasts (representative cells shown by arrows). B. Quantification of transmigrated PBMC from the endothelial channel into the matrix. n>3 devices. C. Quantification of transmigrated Th17 cells from the endothelial channel to the matrix. n>3 devices. P values shown on the graphs.

## Discussion

PSC, like other cholangiopathies, is challenging to model due to its multifactorial etiology, heterogeneity and complex microenvironment (4). Cholangiocyte organoids are used widely for the study of several cholangiopathies. However, using only a single cell type does not capture the interactions between cholangiocytes and different cell types. Vascularized liver organoids have been developed by coculturing iPSC-derived hepatic progenitor cells, mesenchymal cells and endothelial cells to study the inter-lineage interactions during early disease development (29, 30); however, although these organoids formed bile canaliculi, the random organization of the vasculature, the lack of bona-fide bile ducts and inability to perfuse either vessels or ducts has limited their application in the study of cholangiopathies.

Here we demonstrate that endothelial cells, fibroblasts and cholangiocytes can be cocultured in a VBDOC to recapitulate the vascular-biliary interface structurally and functionally. Unlike one tubular model with cholangiocytes that grow on a collagen-coated polyethersulfone hollow fiber membrane with reversed polarity (31, 32), the VBDOC consists of a tubular bile duct within a natural scaffold embedded with mesenchymal cells, adjacent to a 3D vascular vessel that is cultured in endothelial medium. The accessible lumen provides a tractable tool for studying cholangiocyte-bile interactions. Surrounding fibroblasts embedded in a natural and degradable matrix provide the elements for dissecting fibro-obliterative diseases and studying the development of structures such as branches and peribiliary glands.

In addition, the VBDOC is a potential tool to study the role of mechanosensing organelles such as primary cilia. Open endothelial cell and cholangiocyte channels provide the chance to incorporate mechanical stimuli such as flow into the model and thereby study diseases like ciliopathies. Here, by comparing alignment and permeability under static and flow conditions, we demonstrated that endothelial cells respond to shear flow differently from cholangiocytes. The separation between the channels allows collection of medium from each channel for separate analyses and application of distinct mechanical and biochemical stimuli to each, mimicking distinct physiological microenvironments and enabling study of the contribution of mechanical force and biochemical factors to pathology.

Organ-on-a-chip technology is usually limited by cell availability and the need to choose between cell lines with poor fidelity and primary cells in limited supply. Integrating organoid and organ-on-a-chip technology has recently emerged as a superior, synergistic strategy because it enables propagation of primary cells that maintain their original phenotype (14, 33). For example, cholangiocyte organoids can be propagated almost infinitely *in vitro* while still preserving their physiological or pathological features (10, 11, 15). We demonstrated that the VBDOC can be constructed with cholangiocyte organoids from human tissue and bile. These cholangiocytes, when cultured in the device, display a biliary phenotype, with polarity and transport and barrier functions. Moreover, consistent with PSC organoids (11), cholangiocytes in the VBDOC maintained the ability to secret proinflammatory cytokines including CCL20 in response to stimulation by IL-17. By perfusing immune cells through the vascular channel, we modeled the recruitment of immune cells such as PBMC and differentiated Th17 cells to the bile duct. Therefore, the VBDOC could be a valuable platform to study inflammatory and immune-mediated cholangiopathies.

VBDOC devices have limitations due to the nature of the organ-on-a-chip technique. Their fabrication is labor intensive and thus high-throughput experiments are difficult. Additionally, the relatively low number of cells in the device make it hard to perform traditional experiments like Western blotting. Small *in vivo* ducts (several microns) are not possible to replicate because of difficulty generating and seeding the channels. Despite these limitations, the VBDOC provides many benefits over conventional methods and has the potential to be a valuable tool to carry out mechanistic research.

## Author contributions

Y.D. contributed to study concept and design and drafting of the manuscript, and acquisition, analysis and interpretation of data. K.G. assisted in acquisition, analysis and interpretation of data. A.H.-Z., O.W.-Z. contributed human cholangiocyte organoids from tissue, C.S. and J.L.B. provided human cholangiocyte organoids from bile, J.L. participated in cell isolation, C.L. and A.N. contributed human biliary tissue, W.J.P. assisted in the design of the microfluidic device and contributed the algorithm of permeability calculation; R.G.W. conceived ideas and designed the research, provided critical revision of the manuscript, obtained funding and supervised the study.

## Acknowledgments

We gratefully acknowledge the assistance of the following core facilities and individuals at the University of Pennsylvania: the Perelman School of Medicine Cell and Developmental Biology Microscopy Core and Andrea Stout, the Human Immunology Core, and the NIDDK Center for Molecular Studies in Digestive and Liver Diseases Molecular Pathology and Imaging Core (NIH P30 DK050306).

## Abbreviations

PSC: primary sclerosing cholangitis
VBDOC: vascularized bile duct-on-a-chip
CCCM: complete cholangiocyte culture medium
EGM: endothelial growth medium
HUVEC: human umbilical vein endothelial cell
ECM: extracellular matrix
PDMS: polydimethylsiloxane
FITC: fluorescein isothiocyanate
PBS: phosphate buffered saline
Pd: permeability coefficient
PFA: paraformaldehyde
BSA: bovine serum albumin
DAPI: 4′,6-diamidino-2-phenylindole
ZO-1: zona occludens 1
ASBT: apical sodium-dependent bile salt transporter
CFTR: cystic fibrosis transmembrane conductance regulator
SCTR: secretin receptor
ICAM-1: intercellular cell adhesion molecule-1
VCAM-1: vascular cell adhesion molecule-1
LPS: Lipopolysaccharide
PBMC: peripheral blood mononuclear cell

## Figures and Legends

**Figure S1.**
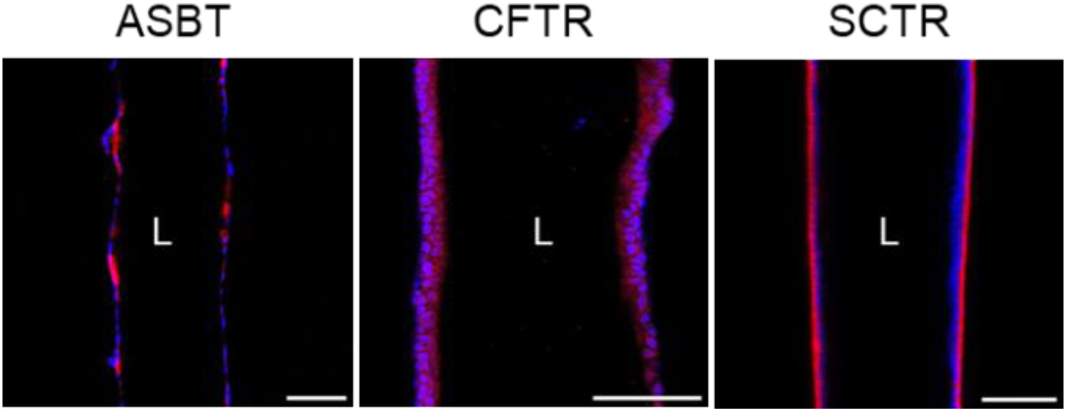
Immunofluorescence images of a biliary channel lined with bile-derived PSC cholangiocytes stained with antibodies against ASBT, CFTR and SCTR (middle cross-sectional view). Nuclei shown by DAPI staining (blue). Scale bars: 100 μm.

**Figure S2.**
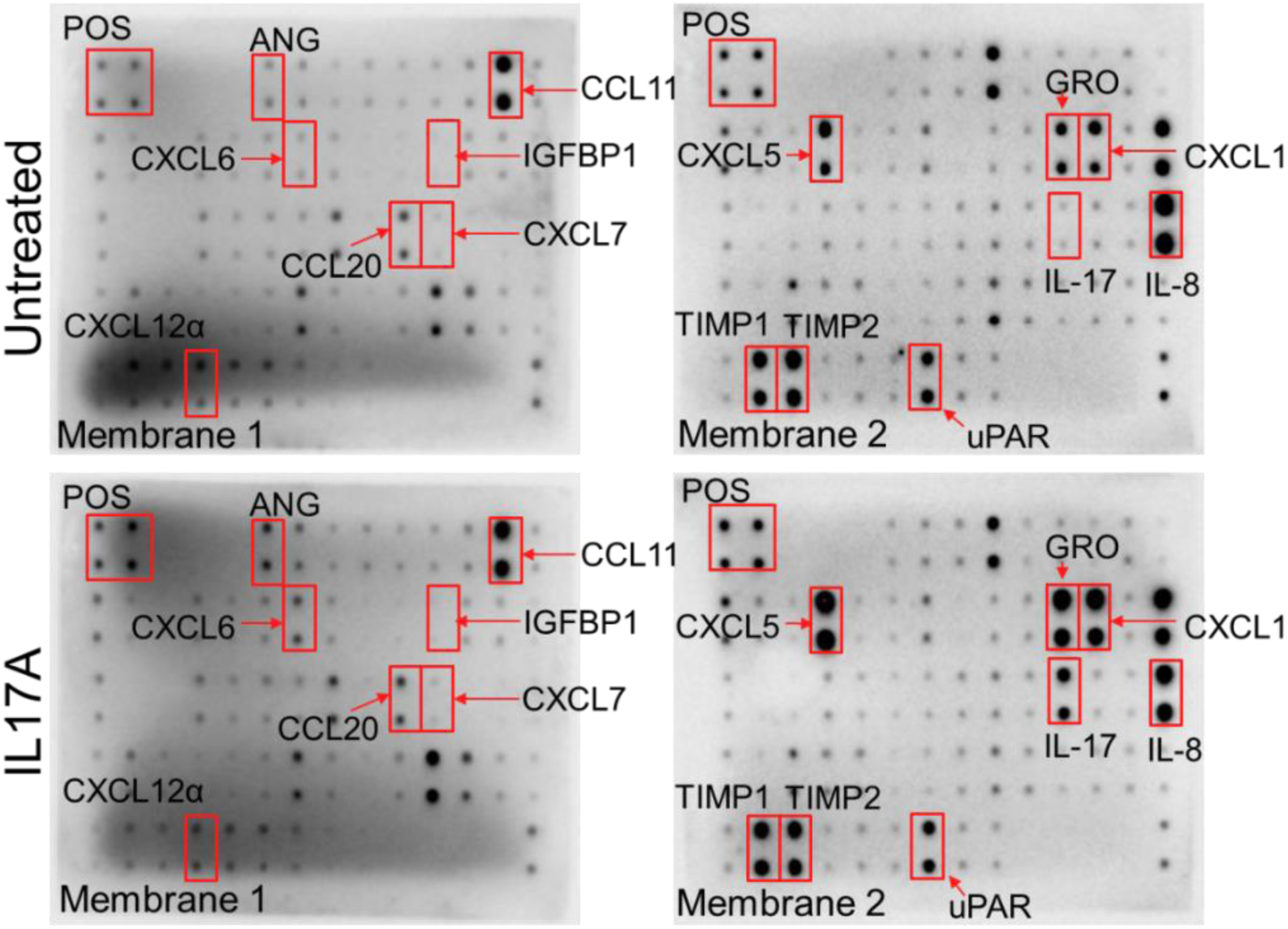
Human cytokine array of supernatants from bile-derived cholangiocytes from PSC patient in the channel under no treatment (upper panels) or treatment with IL-17A (lower panels).

**Figure S3.**
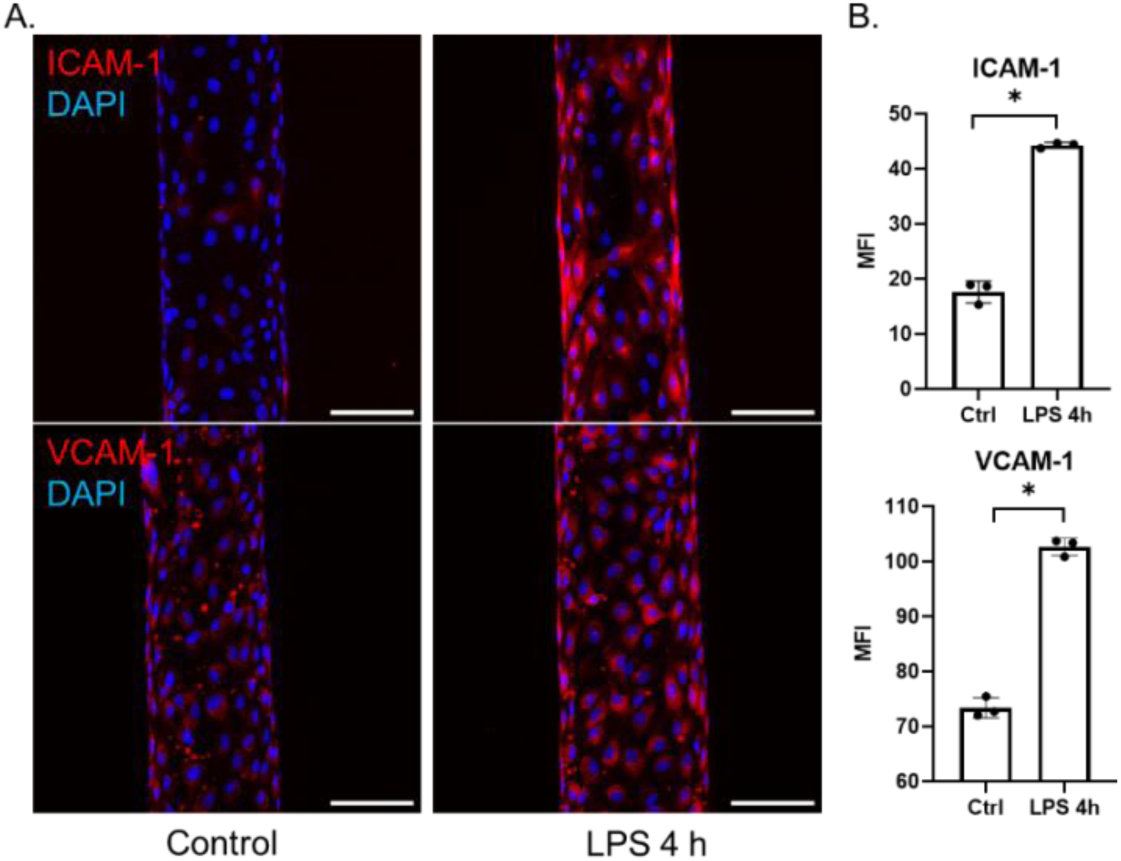
Endothelial cells in channels increased expression of adhesion molecules after LPS stimulation. A. Immunofluorescence images of untreated and LPS-stimulated (4 h) HUVECs stained with antibodies against ICAM-1 (upper row) and VCAM-1 (lower row) (projected top view). Nuclei shown by DAPI staining (blue). Scale bars: 100 μm. B. Quantified mean fluorescent intensity (MFI) of ICAM-1 (upper row) and VCAM-1 (lower row) staining of the HUVEC channel with or without LPS stimulation for 4 h, n=3.

**Figure S4.**
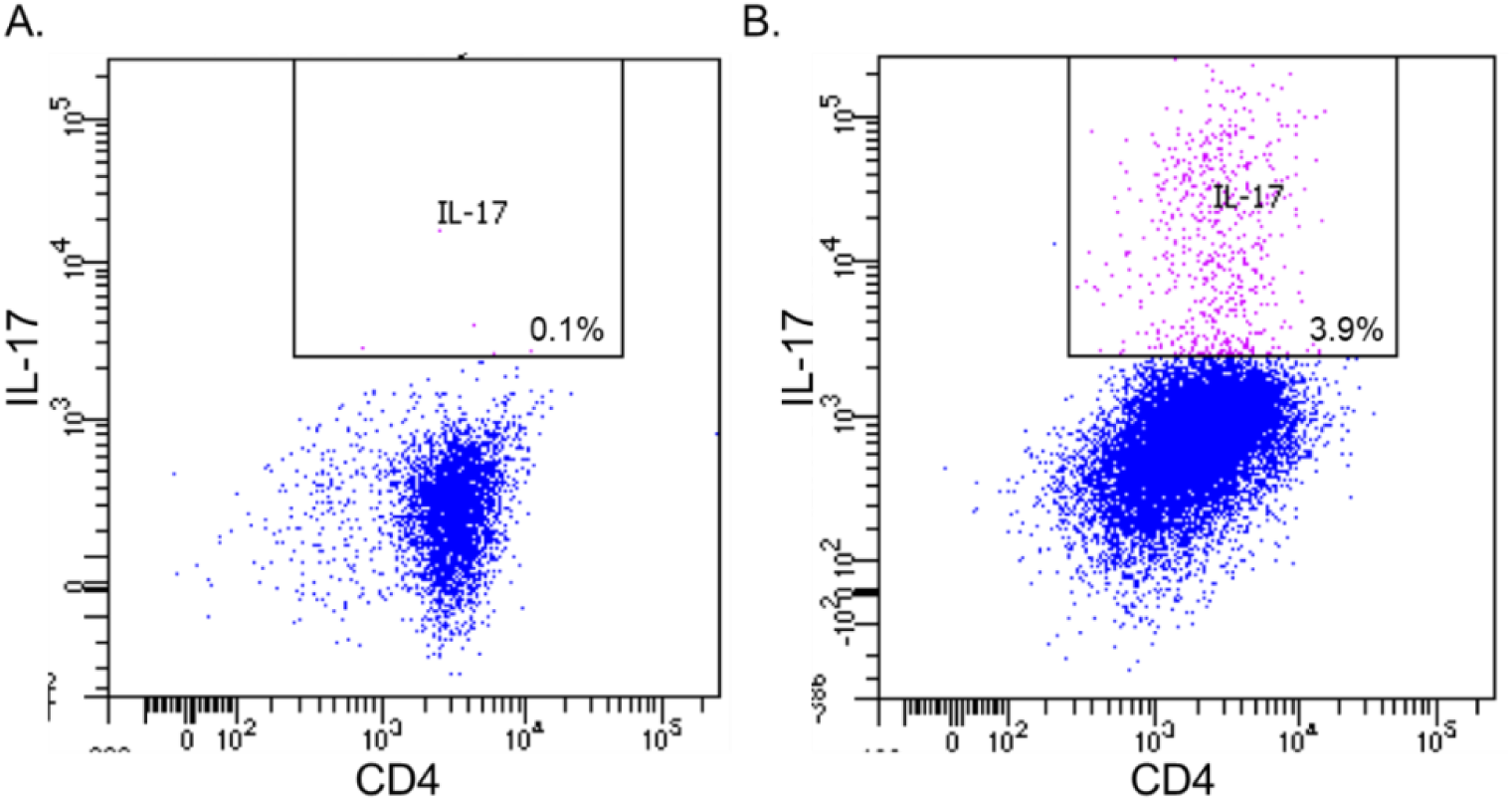
Representative *in vitro* Th17 differentiation from CD4 T cells. A. Representative gating strategy according to intracellular IL-17 staining of CD4+ cells cultured in regular medium. B. Representative gating strategy according to intracellular IL-17 staining of CD4+ cells cultured in differentiation medium.

**Table S1.** Antibodies used.

| Antigen | Concentration/Dilution | Company& Cat. No |
| --- | --- | --- |
| DAPI | 1:5000 | Thermo Fisher Scientific<br>D1306 |
| Phalloidin | 1:100 | Thermo Fisher Scientific<br>R415 |
| K19 | 1:100 | Developmental Studies<br>Hybridoma Bank<br>TROMA-III |
| E-cadherin | 1:100 | Cell Signaling<br>3195S |
| K7 | 1:100 | Abcam<br>ab68459 |
| GGT | 1:100 | Abcam<br>ab55138 |
| EPCAM | 1:100 | Abcam<br>ab239273 |
| Collagen IV | 1:100 | Abcam<br>ab6586 |
| Laminin | 1:100 | Abcam<br>ab11575 |
| Acetylated α-tubulin | 1:100 | Thermo Fisher Scientific<br>32-2700 |
| Apical sodium bile salt<br>transporter (ASBT) | 1:100 | Abcam<br>ab203205 |
| CFTR | 1:100 | Abcam<br>ab2916 |
| SCTR | 1:100 | Abcam<br>ab234830 |
| ZO-1 | 1:100 | Thermo Fisher Scientific<br>61-7300 |
| Occludin | 1:100 | Thermo Fisher Scientific<br>71-1500 |
| Claudin 1 | 1:100 | Thermo Fisher Scientific<br>37-4900 |
| ICAM-1 | 1:100 | Abcam<br>ab214944 |
| VCAM-1 | 1:100 | Abcam<br>ab194319 |
| CD4 | 1:100 | Miltenyi<br>130-120-772 |
| IL-17 | 1:100 | Miltenyi<br>130-120-551 |

**Table S2.** Array map for human cytokine array. A. Membrane 1. B. Membrane 2.

|  |  |  |  |  |  |  |  |  |  |  |  |  |  |
| --- | --- | --- | --- | --- | --- | --- | --- | --- | --- | --- | --- | --- | --- |
| POS | POS | NEG | NEG | BLANK | ANG | BDNF | CXCL13 | BMP4 | BMP6 | CCL23 | CNTF | EGF | CCL11 |
| POS | POS | NEG | NEG | BLANK | ANG | BDNF | CXCL13 | BMP4 | BMP6 | CCL23 | CNTF | EGF | CCL11 |
| CCL24 | CCL26 | FGF-6 | FGF-7 | Flt-3 Ligand | CX3CL1 | CXCL6 | GDNF | GM-CSF | CCL1 | IFN gamma | IGFBP1 | IGFBP2 | IGFBP4 |
| CCL24 | CCL26 | FGF-6 | FGF-7 | Flt-3 Ligand | CX3CL1 | CXCL6 | GDNF | GM-CSF | CCL1 | IFN gamma | IGFBP1 | IGFBP2 | IGFBP4 |
| IGF-1 | IL-10 | IL-13 | IL-15 | IL-16 | IL-1 alpha | IL-1 beta | IL-1ra | IL-2 | IL-3 | IL-4 | IL-5 | IL-6 | IL-7 |
| IGF-1 | IL-10 | IL-13 | IL-15 | IL-16 | IL-1 alpha | IL-1 beta | IL-1ra | IL-2 | IL-3 | IL-4 | IL-5 | IL-6 | IL-7 |
| Leptin | LIGHT | CCL2 | CCL8 | CCL7 | CCL13 | M-CSF | CCL22 | CXCL9 | CCL15 | CCL20 | CXCL7 | NT-3 | CCL18 |
| Leptin | LIGHT | CCL2 | CCL8 | CCL7 | CCL13 | M-CSF | CCL22 | CXCL9 | CCL15 | CCL20 | CXCL7 | NT-3 | CCL18 |
| PDGF BB | CCL5 | SCF | CXCL12 alpha | CCL17 | TGF beta1 | TGF beta3 | TNF alpha | TNF beta | BLANK | BLANK | BLANK | BLANK | POS |
| PDGF BB | CCL5 | SCF | CXCL12 alpha | CCL17 | TGF beta1 | TGF beta3 | TNF alpha | TNF beta | BLANK | BLANK | BLANK | BLANK | POS |

Table S2. Array map for human cytokine array. A. Membrane 1. B. Membrane 2.
|  |  |  |  |  |  |  |  |  |  |  |  |  |  |
| --- | --- | --- | --- | --- | --- | --- | --- | --- | --- | --- | --- | --- | --- |
| POS | POS | NEG | NEG | BLANK | Acrp30 | AgRP | ANGPT2 | AREG | Axl | bFGF | b-NGF | BTC | CCL28 |
| POS | POS | NEG | NEG | BLANK | Acrp30 | AgRP | ANGPT2 | AREG | Axl | bFGF | b-NGF | BTC | CCL28 |
| CTACK | Dtk | EGFR | CXCL5 | Fas | FGF-4 | FGF-9 | G-CSF | GITR Ligand | GITR | GRO | CXCL1 | CCL16 | HGF |
| CTACK | Dtk | EGFR | CXCL5 | Fas | FGF-4 | FGF-9 | G-CSF | GITR Ligand | GITR | GRO | CXCL1 | CCL16 | HGF |
| ICAM-1 | ICAM-3 | IGFBP3 | IGFBP6 | IGF-1 sR | IL-1R4 | IL-1 R1 | IL-11 | IL-12 p40 | IL-12 p70 | IL-17 | IL-2R alpha | IL-6R | IL-8 |
| ICAM-1 | ICAM-3 | IGFBP3 | IGFBP6 | IGF-1 sR | IL-1R4 | IL-1 R1 | IL-11 | IL-12 p40 | IL-12 p70 | IL-17 | IL-2R alpha | IL-6R | IL-8 |
| CXCL11 | XCL1 | MIF | CCL3 | CCL4 | CCL19 | MSP alpha | NT-4 | OPG | OSM | PLGF | sfp130 | sTNFRII | sTNFRI |
| CXCL11 | XCL1 | MIF | CCL3 | CCL4 | CCL19 | MSP alpha | NT-4 | OPG | OSM | PLGF | sfp130 | sTNFRII | sTNFRI |
| CCL25 | TIMP-1 | TIMP-2 | THPO | TRAIL R3 | TRAIL R4 | uPAR | VEGF | VEGF-D | BLANK | BLANK | BLANK | BLANK | POS |
| CCL25 | TIMP-1 | TIMP-2 | THPO | TRAIL R3 | TRAIL R4 | uPAR | VEGF | VEGF-D | BLANK | BLANK | BLANK | BLANK | POS |

